# Diverse but temporally stable virome in a 1.2 km deep karst aquifer accessed via Moab Khotsong mine, South Africa

**DOI:** 10.1101/2025.10.27.684889

**Authors:** Nathaniel E. Jobe, Daniel S. Jones, Julio César Castillo Hernandez, Devan M. Nisson, Cassandra H. Skaar, Tullis C. Onstott, Thomas L. Kieft

## Abstract

Despite their ubiquity, viruses remain understudied in many of Earth’s environments. The deep subsurface contains a large portion of Earth’s biomass and can be expected to hold a rich store of viruses as well. We investigated the viral community in a 1.2-kilometer-deep dolomitic aquifer accessed in the Moab Khotsong gold mine in South Africa by microscopically quantifying virus-like particles (VLP) and cells and conducting a metagenomic survey of viruses and their associations with the microbial community. VLP:cell ratios were consistently lower (∼1:1) than those found in most shallower aquifers. Viral sequences were recovered from both the <0.2 µm fraction, presumably representing free virions, and the >0.2 µm fraction that also included microbial cells. Viruses were diverse and 78% were novel, and the viral communities were relatively unchanged at four sampling dates spanning five years. Host prediction indicated viral infection of the dominant microbes, including *Thiomicrospiraceae* and *Rhodocyclaceae*, with temperate phages. Viral genomes include auxiliary metabolic genes (AMGs), with most of them being Type II, which favor host persistence. Taken together, VLP:cell ratios, type II AMGs, and temporally stable viromes point to dominance of lysogeny and piggyback-the-persistent viral-host interactions in the deep aquifer. The many novel viruses observed here suggest that the terrestrial subsurface is an untapped reservoir of viral diversity.

## Introduction

The deep subsurface contains a significant proportion of Earth’s estimated biomass [1]. It can be broadly separated into marine and terrestrial realms, the latter being operationally defined as one or more meters below land surface (mbls) [2]. Deep terrestrial subsurface life functions without the direct influence of photosynthesis [3] and has an estimated abundance of 2 × 10^29^ to 6 × 10^29^ cells and species richness of ∼10^12^ [4]. Comprising ∼85% of Earth’s bacteria and archaea [1], these ecosystems are home to metabolically diverse communities involved in global biogeochemical cycling [3, 5]. However, due to its vast size [4, 5] and limited accessibility of sampling [5], the deep terrestrial subsurface represents one of the least studied environments on Earth.

Viruses are ubiquitous and drive important microbial community interactions [6, 7], the full extent of which remains unknown in most systems. Viral interactions with microbial communities can generally be separated according lytic and lysogenic cycles. The lytic viral life cycle, resulting in the lysis of the host, is often associated with “kill-the-winner” dynamics and the viral shunt [8, 9], which can significantly impact microbial biomass and nutrient turnover. Kill-the-winner describes a process in which viruses infect fast-growing or dominant hosts, leading to fluctuating “boom-and-bust” population patterns for both host and virus [9]. The viral shunt refers to viral lysis of host cells releasing organic molecules that can be taken up and metabolized by other cells [10, 11]. The lysogenic cycle, in which the viral genome replicates within the host as an integrated or extrachromosomal prophage, is associated with “piggyback-the-winner” or “piggyback-the-persistent” dynamics [9, 12] and more stable host populations. Viruses can also impact hosts through accessory metabolic genes (AMGs), which are host-like genes that are associated with altering host metabolism to effect a more successful infection [13]. Class I AMGs encode metabolic pathways like catabolism [14, 15] to enhance energy production for virus production in the lytic cycle [15]. Class II AMGs play peripheral roles in metabolism and are associated with physiological regulation and maintaining mutualism [13, 14] during lysogeny.

Previous research on viruses in aquifers has mostly focused on exogenous pathogenic viral strains [16, 17] and their transport through aquifers [18]. Few studies have examined naturally occurring viral communities in aquifers; these have included quantification of virus-like particles (VLPs) [12, 19–22], metagenomic sequencing [23–27], culturing [28], single-cell sequencing [29] and transcriptomic analyses [24, 27]. There are even fewer studies that have characterized endogenous viromes of groundwater systems; those available report that changes in fluid geochemistry and microbial community compositions lead to concomitant shifts in associated viral communities [19, 24]. Studies at the Äspö Hard Rock Laboratory in Sweden represent some of the deepest viral studies to date, ranging from 65 to 450 mbls [21, 27, 28]. Other groundwater studies have shown that changes in geochemistry and associated changes in microbial communities lead to concomitant shifts in the viral communities [19, 24]. Variations in AMG profiles associated with lytic or temperate viruses have also been found in deep subsurface sites with class I AMGs dominant in groundwater samples in China [25] and class II AMGs appearing to be more dominant in aquifers in New Zealand [24] and the Äspö Hard Rock Laboratory [27].

The goal of this study was to analyze the viral community of a deep (1200 mbls) dolomitic karst aquifer in the Witwatersrand Basin in South Africa, providing the first virome characterization of a groundwater aquifer deeper than 1 kbls. This aquifer, accessed in the Moab Khotsong gold mine in South Africa, has previously been studied for its groundwater geochemistry [30, 31] and microbiology [32]. We quantified VLPs and cells, performed metagenomic analyses of viruses and microbes, and used viral sequence data to determine taxonomic classification, host prediction, and genetic capabilities, including AMGs. As the VLP:cell ratio in other deep subsurface sites tends to be lower than the average of 10 that is common in the surface ocean [7, 33, 34], we hypothesized that VLP:cell ratios would be <10. Additionally, based on the remote and unusual nature of the aquifer habitat, we hypothesized that most viral sequences would be novel. We further hypothesized that the most common viruses would be associated with abundant hosts, and would be consistent with either kill-or piggyback-the-winner interactions, based on the prevalence of these phage-host dynamics in other systems [8, 9] and depending whether the deep aquifer community undergoes dynamic fluctuations over multi-year timescales.

## Materials and methods

### Aquifer characteristics and sampling methods

Samples were collected at the 1200 level of Moab Khotsong mine at 1200 mbls by attaching a sterilized stainless-steel manifold to a rubber hose extending from the dam of the main pump station at that level. The aquifer accessed at the 1200 level represents a paleo-meteoric fluid from a karstic aquifer in the 2.2-2.5 Ga Transvaal dolomites [35]. Samples for chemical analysis collected in sterilized, N_2_-flushed serum vials and refrigerated until analyzed. Two 10-L water samples were collected and filtered on site using a 0.2-µm Whatman AS 36 Polycap filter, with the filtrate intended for viromic analyses (<0.2 µm fraction) and the filter intended for analysis of the microbial community and any viruses retained on the filter (>0.2 µm fraction). Additional 0.2-µm Millipore Sterivex filters were used to collect cells. Samples for cell and VLP counting (unfiltered and 0.2 µm-filtered, respectively) were collected in 50-mL conical centrifuge tubes, and fixed on site to a final concentration of 3.7% formaldehyde (2023) or paraformaldehyde (2024). Samples were transported to a surface laboratory, and then frozen at –20 °C (filters) or refrigerated at 4 ℃ (filtrate, water and fixed samples). Samples were transported from South Africa to New Mexico Institute of Mining and Technology on frozen ice packs (“blue ice”) and then refrigerated (water) or frozen at –80 ℃ (filters).

### Site geochemistry

Sulfide, iron (ferric and total) and dissolved oxygen were measured on-site using Chemet kits (CHEMetrics, Midland, VA, USA). Temperature, pH, ORP, EC and TDS were measured with a HANNA pHep and ORP probe. The pH was measured using Environmental Protection Agency (EPA) method 150.1. Cations and anions were measured using inductively coupled plasma optical emission spectroscopy (ICP-OES) using EPA method 200.7 for cations and 300.0 for anions. Trace metals were measured via inductively coupled plasma mass spectrometry (ICP-MS) using EPA method 200.8. Alkalinity was measured by titration (standard method SM 20320 B [36]) and hardness was calculated by standard method SM 340B [36].

### Cell and VLP counting

Both filtered and unfiltered fixed samples were used for cell and VLP counts. Sample were diluted in 0.2-µm filtered, autoclaved MilliQ water and combined with 20× SYBR Gold [37]. Samples were then filtered through a 0.02-µm Whatman Anodisc membrane and then rinsed with sterile water to remove unbound SYBR Gold. The Anodisc membrane was mounted onto a slide using VECTASHIELD PLUS antifade mounting medium (Vector Laboratories). Cells and VLPs were counted at 1000× magnification using a BX63F epifluorescence microscope (Olympus, Tokyo, Japan) with an ORCA-Spark camera (Hamamatsu, Japan) running Olympus cellSens dimensions software 3.1.1.

### Microbial DNA extraction and analysis

Polycap and Sterivex filters were aseptically dissected and the DNA was extracted from the filter material using DNeasy PowerSoil Pro (2023) or PowerWater (2024) kits (Qiagen) according to the manufacturer’s protocol except that samples were vortexed for 5, 10, and 15 minutes and then recombined. Resulting DNA was quantified using Nanodrop spectrophotometer (Thermo-Fisher Scientific) and Qubit fluorometer (high sensitivity DNA kit; Invitrogen). Metagenomic libraries of microbial communities (>0.2 μm fraction) were prepared and sequenced at the University of Minnesota Genomics Center using an Illumina NovaSeq (paired end, 2 × 150 bp) for samples collected in 2022 and an Element Biosciences AVITI platform (paired end, 2 × 150 bp) for samples from 2024. rRNA gene libraries were prepared by amplifying the V4 hypervariable region of the 16S rRNA gene using 515f/806r primers [38] as in Jones et al. [39]. Complete details on the preparation and analysis of metagenomic and rRNA gene libraries are provided in the supplementary information.

### Viral concentration, DNA extraction, and sequencing

Viruses were concentrated from the filtered samples by two methods: iron flocculation [40] and tangential flow filtration (TFF) [41]. The iron flocculation method involved sorption of viruses to iron-oxide precipitate (formed by adding FeCl_3_•6H_2_O), which was then captured on 0.8-µm pore size polycarbonate filters and stored at 4 ℃. The iron oxide was then dissolved, releasing viruses using a 0.1 M EDTA-0.2 M MgCl_2_-0.2 M oxalic acid buffer (pH 6). TFF was performed using a Biomax 100-kDa polyethersulfone membrane cassette and a Millipore Lab Scale TFF system.

The resulting 50 mL concentrate was stored at 4 ℃. Volumes of the iron flocculation resuspension and the TFF retentate were reduced to 4 mL using Millipore Amicon 100-kDa centrifugal filter tubes and then to 30 µL using Nanosep microcentrifuge filters. Final viral suspensions were treated with DNase I following the method of Hurwitz et al. [42] to remove free DNA. DNA was extracted from DNase-treated samples using Wizard DNA Purification Resin and Wizard Mini Columns. Extracted viral DNA was sequenced at the University of Minnesota Genomics Center using an Illumina NovaSeq or Element Biosciences AVITI (paired end, 2 × 150 bp).

### Viral sequence identification, gene annotation, and host-network analyses

Viral sequences were identified using VirSorter2 v2.2.4 [43], DeepVirFinder v1.0 [44], and VIBRANT v1.2.1 [45]. Gene-sharing network analysis was performed on the high-quality contigs and bins using vConTACT2 v 2.6.3 [46, 47]. For taxonomic assignment, vConTACT2 was used in conjunction with files from INPHARED [48]. The output network file and genome-by-genome overview file were then used with graphanalyzer.py v1.6.0, a script made for the MetaPhage pipeline [49], to assign taxonomy to the clustered high-quality viral sequences according to ICTV taxonomy. Taxonomy information from graphanalyzer was visualized using Krona plots [50]. Viruses classified to the family *Microviridae* were removed from downstream analysis due to the ΦX174 sequencing control in the samples. An additional network was created using sequences from IMGVR v4.1 [51] to identify the similarity of these viral sequences to uncultured viruses. For this, viruses associated with marine aquifers, deep subsurface aquifers, deep subsurface groundwater, and freshwater groundwater were extracted from the nucleotide high confidence release. Host prediction of the high-quality contigs was made using iPHoP v1.3.3 [52]. AMGs from the co-assembled viruses were identified using a combination of VIBRANT and DRAM-v. Full viral gene annotation was performed using pharokka v1.7.3 [53] and phold v0.2.0 [54]. Complete details of metagenomic analysis of the >0.2 μm and <0.2 μm fractions are provided in the Supplementary Material.

## Results

### Aquifer geochemistry

Geochemical parameters varied slightly across the three main collection periods (Table S1). The aquifer was alkaline with a pH between 7.9 and 8.6 and had total dissolved solids between 1029 and 1086 mg/L, in the range of brackish water. Across the three sampling dates, trace sulfide and dissolved oxygen concentrations were detectable, while ferrous iron and total iron were detectable only in 2022. Nitrate and nitrite were not detected during this sampling but had been detected previously [32]. The alkalinity of the water was mainly due to CaCO_3_ and HCO_3_^-^. The major ions with the highest concentrations were sodium, chloride, and sulfate.

### Cell and VLP counts

Cell counts averaged 1.1 × 10^6^ cells/mL (standard deviation: 3.8 × 10^5^ cells/mL); multiple cell morphologies were observed, including bacilli, coccobacilli, cocci, and spirilla. VLP counts averaged 1.2 × 10^6^ VLP/mL (standard deviation: 2.3 × 10^5^ VLP/ml) for a VLP:cell ratio of ∼1.1.

### 16S rRNA gene amplicon libraries

To characterize the microbial community over time, six amplicon libraries were prepared from samples from filters collected during 2022, 2023, and 2024 (Table S2). Based on the rRNA gene libraries, the most abundant OTUs were as *Sulfuritalea*, *Thioclava*, *Magnetospirillum*, *Thiobacillus*, *Lentimicrobium*, and *Rhodobacter* (Fig. S1). Archaeal sequences represented ≤0.2% of the total sequences. Based on the cluster analysis, samples from the same year cluster together.

### Metagenomic and metaviromic libraries

Metagenomic sequencing was performed on viral concentrates (<0.2 µm fraction) and filters (>0.2 µm fraction, comprising cells along with viruses trapped on the filters) (Tables 1, S3). Metagenomic libraries from the <0.2 um fraction ranged from 7.1 to 18.7 Gb, while libraries from the >0.2 um fraction ranged from 17.2 to 31.6 Gb. A library generated from the 2024 Fe flocculation extract was only 0.2 Gb, and was excluded from subsequent analyses. Each library was sampled individually, and then the five metagenomes and all eight datasets were co-assembled (Table 2). Individual metagenome assemblies from the >0.2 µM fraction had N50 values between 2,348 and 25,877, with 1,439-2,417 and 75-175 contigs >10 kb and >100 kb, respectively. The co-assembly of the five metagenomes had an N50 value of 2,355, with 7,020 and 341 contigs longer than 10 kb and 100 kb, respectively. The co-assembly of the eight datasets had an N50 value of 2,662, with 9,442 and 433 contigs longer than 10 kb and 100 kb, respectively.

**Table 1.** Samples used for viromic analyses.

|  | Viromes<br>(<0.2 µm fraction) |  |  |  | Mixed metagenomes (cells and viruses)<br>(>0.2 µm fraction) |  |  |  |  |
| --- | --- | --- | --- | --- | --- | --- | --- | --- | --- |
| Year | 2023 |  | 2024 |  | 2019 | 2022 |  | 2024 |  |
| Sample name | 100kDa<br>TFF | PC_Fe | 1200L-TFF | FE_clean up | Nisson<sup>a</sup> | BF1 | BF2 | PC_b | Steri_a |
| Sample type | TFF | Iron Flocculation | TFF | Iron Flocculation | Polycarbonate | Polycap | Polycap | Polycap | Sterivex |
| Meta-genome size (Gbp) | 7.126 | 12.869 | 18.650 | 0.974 | 19.749 | 25.197 | 31.589 | 22.341 | 17.168 |

<sup>a</sup>Metagenome dataset from Nisson et al. [32] downloaded from the NCBI Sequence Read Archive (SRA)

**Table 2.** Assembly statistics for the co-assemblies.

|  | Metagenome<br>Co-assembly | Virome/Metagenome<br>Co-assembly |
| --- | --- | --- |
| Number of scaffolds x 10<sup>3</sup> | 483.541 | 588.794 |
| Total size of scaffolds (Mbp) | 762.892271 | 977.862699 |
| Longest scaffold (kbp) | 1183.93 | 1183.73 |
| Mean scaffold size (bp) | 1578 | 1661 |
| Median scaffold size (bp) | 766 | 783 |
| N50 (bp) | 2355 | 2662 |
| L50 (kbp) | 45833 | 52808 |

### Metagenome-assembled genomes (MAGs)

Metagenome-assembled genomes (MAGs) were generated from the co-assembled filter metagenomes (>0.2 µm fraction) to characterize the genomic potential of microbial community members and improve host prediction of viral sequences. A total of 145 bins were generated, of which 85 were >50% complete with <5% contamination as determined by CheckM or CheckM2. These 85 were retained as high-quality bins (Table S4). Of these, 84 represent bacterial bins with only one archaeal bin. Taxonomic classification of the bins based on the Genome Taxonomy Database (GTDB) shows that the majority of bins were classified to the phyla *Pseudomonadota* (n=44), most of which were in class *Alphaproteobacteria* (n=17) and *Gammaproteobacteria* (n=16). Other phyla were *Bacteroidota* (n=10), *Patescibacteria* (n=8), *Desulfobacterota* (n=4), *Bdellovibrionota* (n=2), *Campylobacterota* (n=2), *Actinomycetota* (n=2), *Chloroflexota* (n=2), *Planctomycetota* (n=2), *Bacillota*_I (n=1), *Bipolaricaulota* (n=1), *Krumholzibacteriota* (n=1), *Zixibacteria* (n=1), *Dependentiae* (n=1), *Verrucomicrobiota* (n=1), *Myxcoccota* (n=1) and *Bacillota* (n=1). The only archaeal bin was classified to the genus *Forterrea* in the phylum *Iainarchaeota.* This distribution of phyla is similar to those identified from the 18 MAGs recovered by Nisson et al. [32]. Annotation using DRAM-identified genes related to catabolism, specifically electron transport, hydrogenases, and one-carbon metabolism (Fig. S2). Type I, II, or III CRISPR systems and two Cas proteins were found in 33 of the bins (Fig. S3).

### Virome recovery

In total, 7,950 viral sequences were predicted from the co-assembly (Fig. S4). Following viral binning, a total of 6,286 high-quality viral sequences were generated (488 bins and 5,798 contigs). Based on CheckV quality summaries, 24 are complete viral sequences, 77 are high-quality sequences, 128 are medium-quality, 3,706 are low-quality, and the remaining 2,351 were not determined. Two-way cluster analysis of viral sequences using the actual coverage for the viral contigs and an average coverage of contigs in viral bins revealed that viruses concentrated by iron flocculation or TFF and collected on different dates were very similar to each other but differed from viruses that were captured along with cells on 0.2-µm filters (Fig. 1).

**Figure 1.**
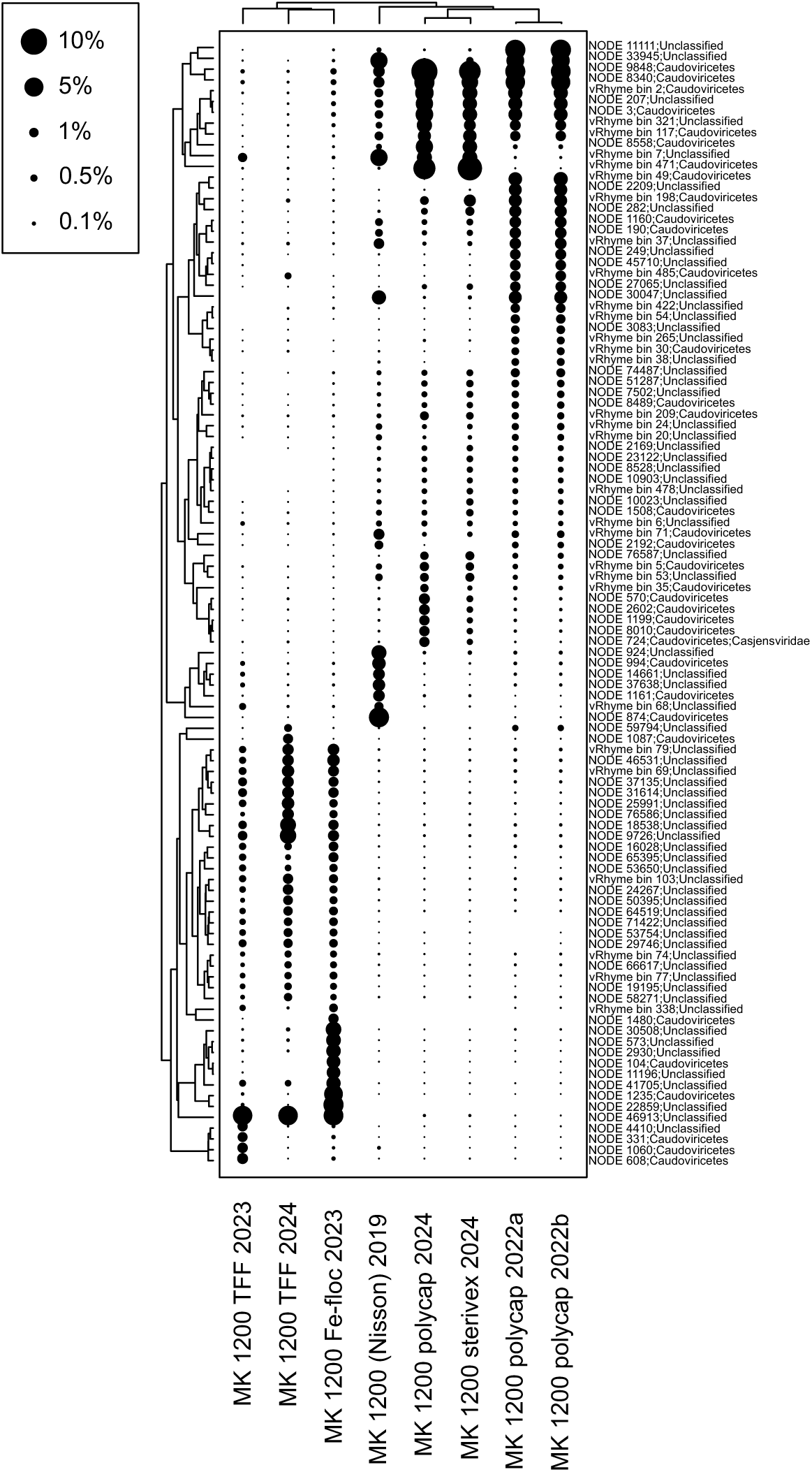
Two-way hierarchical agglomerative cluster analysis of viral sequences, based on coverage. The R-mode cluster analysis (clustering of viral contigs or bins) was calculated using the 100 most abundant predicted viruses, while the Q-mode cluster analyses was calculated using all predicted viruses.

### Taxonomic clustering

Gene sharing network analysis of the viral sequences with the INPHARED database of cultured viruses using vConTACT2 (Fig. S5) shows the relationships of 1,017 sequences in this study to each other and to database viruses. (The remaining 5,269 viruses did not cluster due to no or low homology with viral sequences either in the database or from this study.) Of the 1,017 sequences that clustered, the majority of them were not assigned any taxonomic information (Fig. 2A). The majority of the classified viruses belong to three viral classes: *Caudoviricetes* (n=622), *Faserviricetes* (n=7), and *Tectiliviricetes* (n=1). Of these, 69 were classified to the family level or lower, representing 19 viral families and an additional four viral families without an associated family designation (Fig. 2B). The majority of these families fall within the *Caudoviricetes* class with one family each represented for the other two classes: the *Inoviridae* under the *Faserviricetes* and the *Tectiviridae* under *Tectiliviricetes*. The five most common viral families were classified as *Mesyanzhinovviridae* (n=8), *Schiroviridae* (n=8)*, Winoviridae* (n=7), *Autographiviridae* (n=5), and *Inoviridae* (n=5). An additional network analysis that included viruses not only from RefSeq but also other groundwater metaviromes from IMG/VR showed that 1,367 viral sequences (∼22% of the total number of 1200-level viruses) appeared in a cluster containing at least one IMG/VR or RefSeq virus, leaving ∼78% of 1200-level viruses unrelated to any known viruses (Fig. S6).

**Figure 2.**
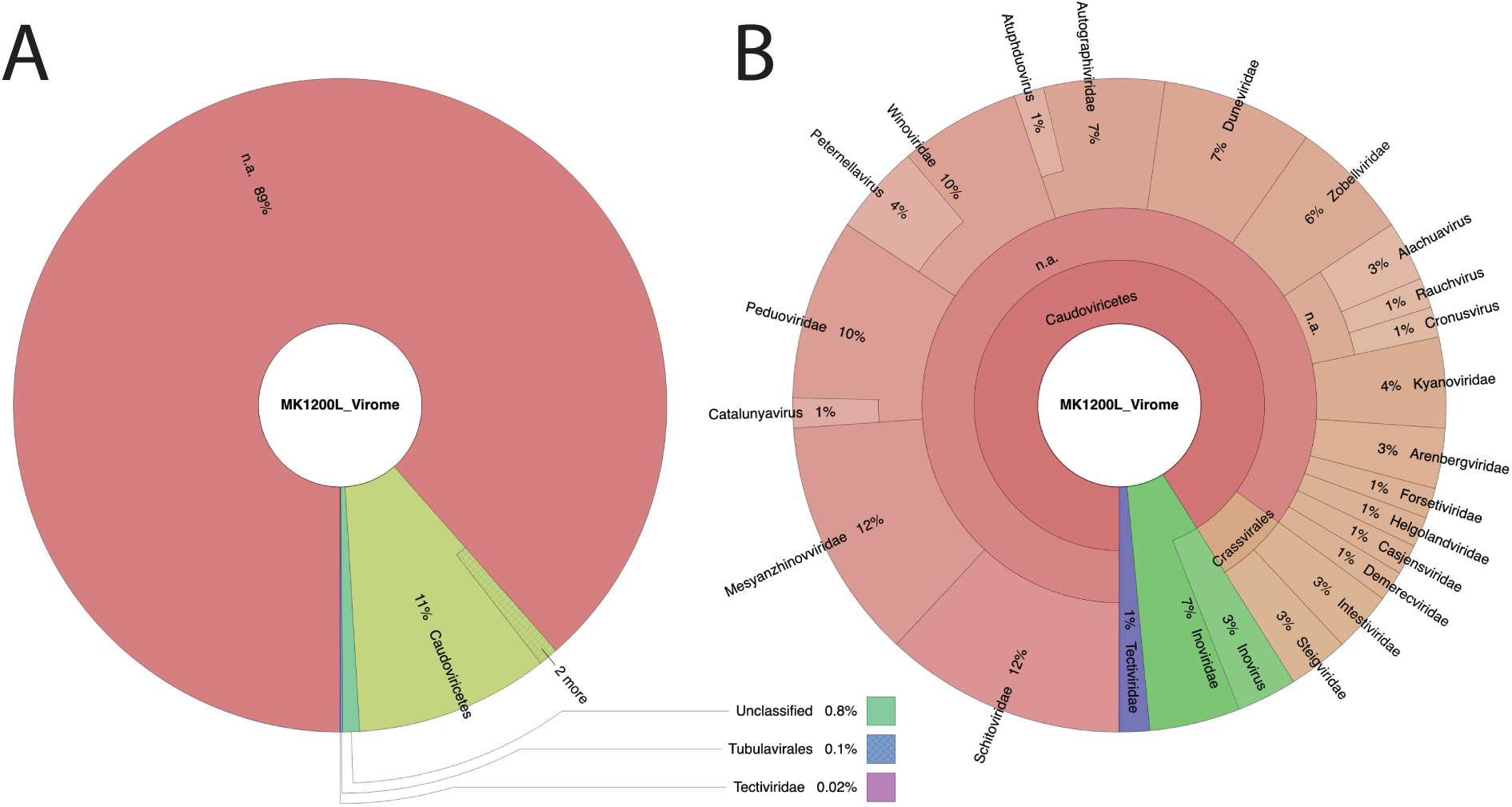
A. Krona plot of all the viral sequences from the coassembly with the taxonomic assignments generated by graphanalyzer and vConTACT2. B. Krona plot of the 69 contigs with a family or lower taxonomic assignment.

We annotated several viruses that were taxonomically classified with vConTACT2 and graphanalyzer (Fig. S7) to compare genome structure and organization to known viruses. Figure S7A presents a viral bin that was classified to the *Crevaviridae* family, a group of uncultured but abundant phages called CrAssphages, which have been found in diverse environments [55,56]. Figure S7B shows the one viral sequence classified to the *Gammatectivirus* genus, which only has one cultured representative [57]; the viral contig includes DNA polymerase characteristic of *Tectiviridae* [58]. Figure S7C presents a sequence classified as to the genus *Inovirus*. These are ssDNA filamentous viruses that are capable of establishing a chronic infection. Members of the *Inoviridae* family are present globally across biomes [59]. Figure S7D presents a viral bin that was classified to the family *Kyanoviridae* members of which contain many AMGs [60].

### Host prediction

Host prediction with iPHoP associated 710 viral sequences with a host from the iPHoP; these represent 300 bacterial genera. No archaeal hosts were predicted for the viral sequences. The most commonly predicted host families include *Moraxellaceae* (n=76), *Burkholderiaceae_*B (n=65), *Rhodobacteraceae* (n=52), *Pseudomonadaceae* (n=48), and *Rhodocyclaceae* (n=39) based on GTDB taxonomy. Focusing on the MAGs added to the iPHoP database from the mixed metagenome coassembly, 37 of the 50 bacterial MAG families had associated viral sequences (Fig. 3). Of the families of the added MAGs, *Rhodobacteraceae* (n=16), *Thiotrichaceae* (n=12), *Thiobacillaceae* (n=10), *Rhodocyclaceae* (n=9), UBA11246 (n=9) and *Thiomicrospiraceae* (n=6).

**Figure 3.**
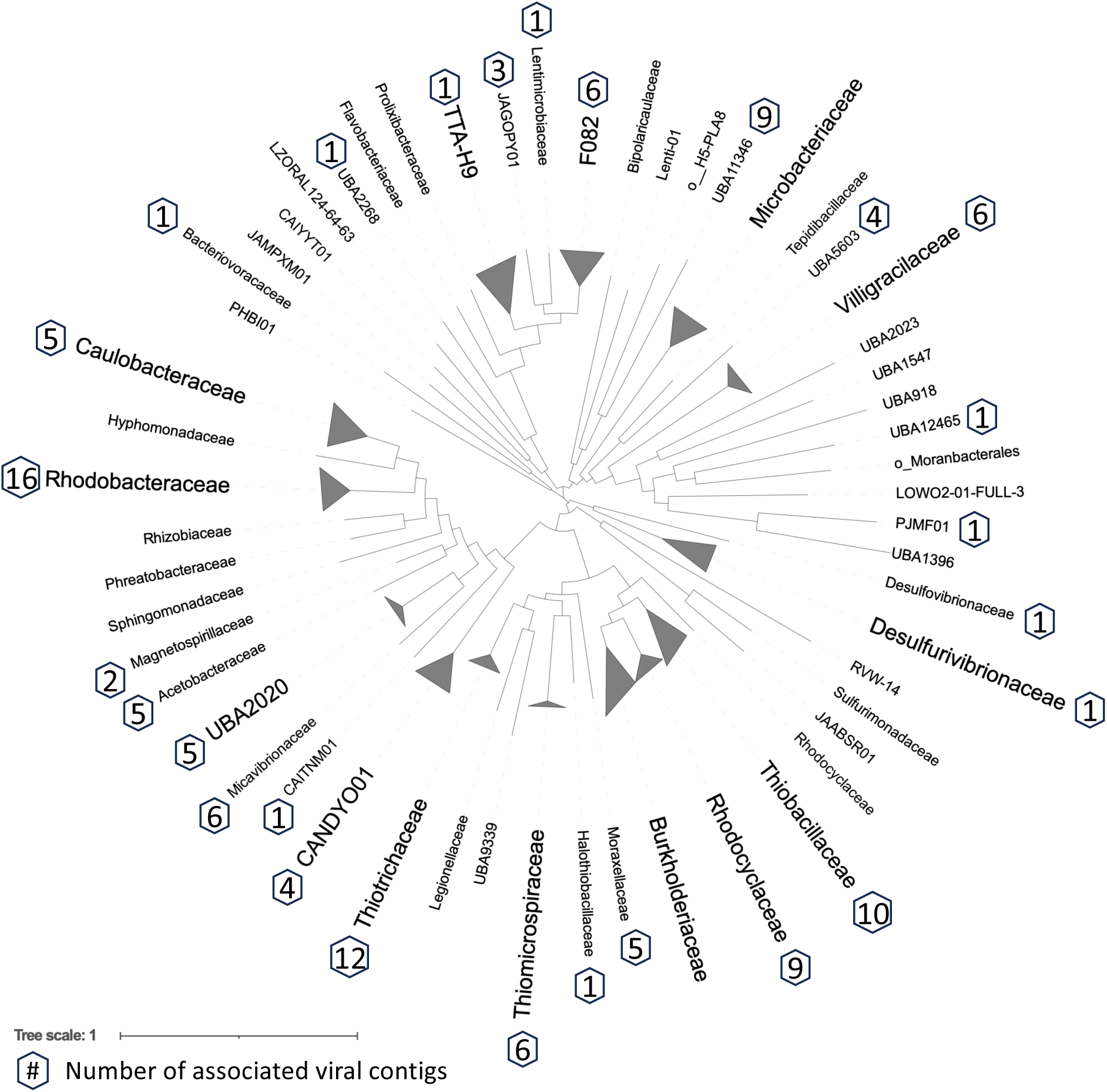
Tree of high-quality metagenome-assembled genome bins based on the GTDB alignment of 120 concatenated phylogenetically informative proteins generated with FastTree and visualized in iTOL. Tips labeled with the family classification, where the bins with the same family-level classification were collapsed. Families with viruses predicted to infect them are signified with a hexagon containing the number of associated viral sequences. Scale bar represents average substitution per site.

### AMGs

AMGs annotated with both VIBRANT and DRAM-v were combined to identify genes related to a variety of metabolic functions (Fig. 4). The most abundant AMGs were related to the metabolism of cofactors, vitamins, and amino acids. Two abundant genes identified as AMGs were one encoding DNA (cytosine-5)-methyltransferase 1 (n=89) involved in DNA methylation and *nrdA*/*nrdB* encoding ribonucleotide reductase (n=19); however, nucleotide metabolism genes were eliminated from further analysis, as they are unlikely to be actual AMGs [14]. The most abundant amino acid metabolism gene was *cysH* (n=19) encoding phosphoadenosine phosphosulfate reductase. The highest number of AMGs within a viral sequence was nine. Seventy-five AMGs were grouped as “Other”, which encode proteins from miscellaneous pathways (Table S5). Class II AMGs outnumbered Class I AMGs 318 to 205.

**Figure 4.**
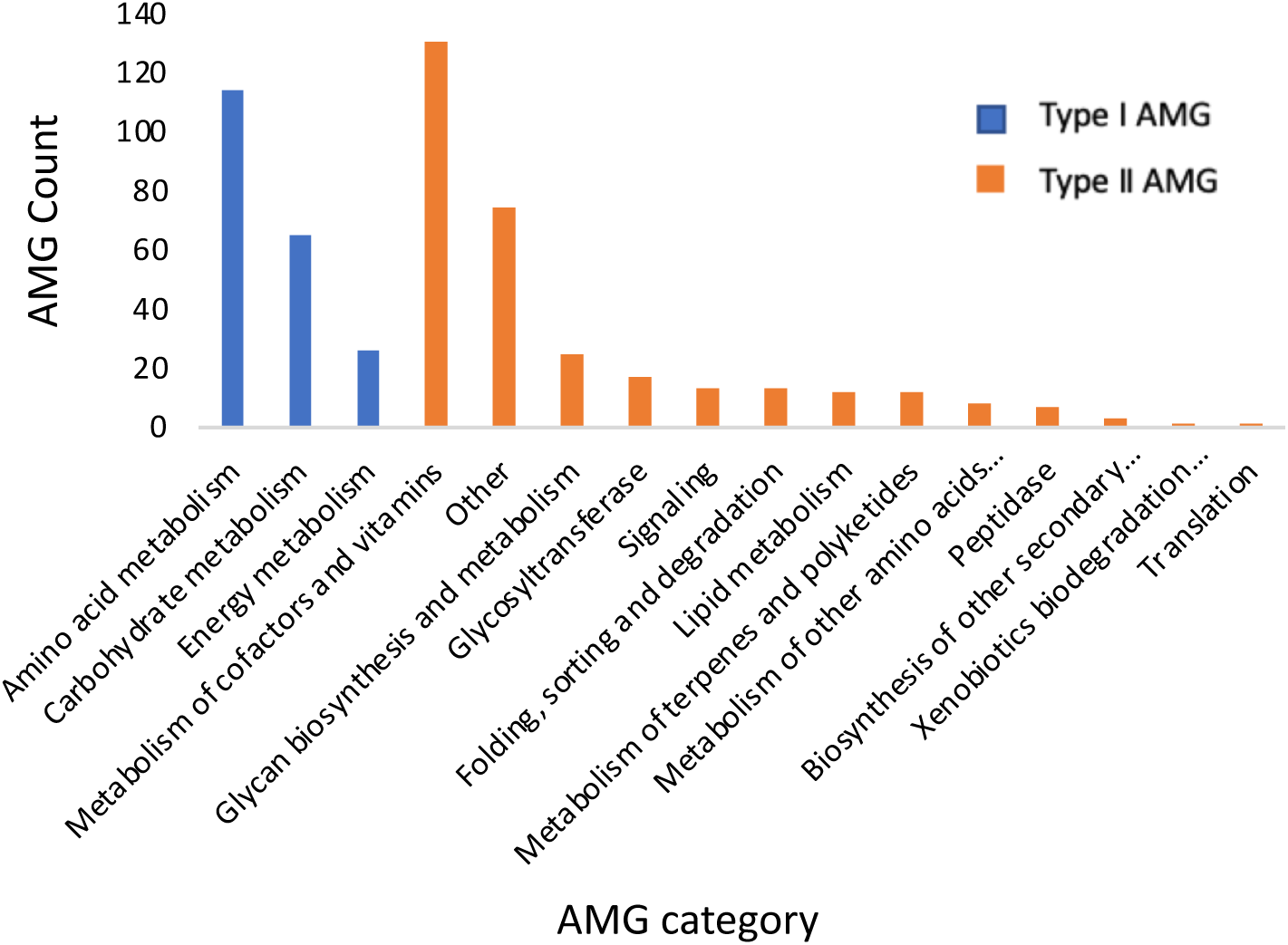
AMGs present in the coassembly. “Other” refers to AMGs encoding proteins from miscellaneous pathways (Table S5).

**Figure 5.**
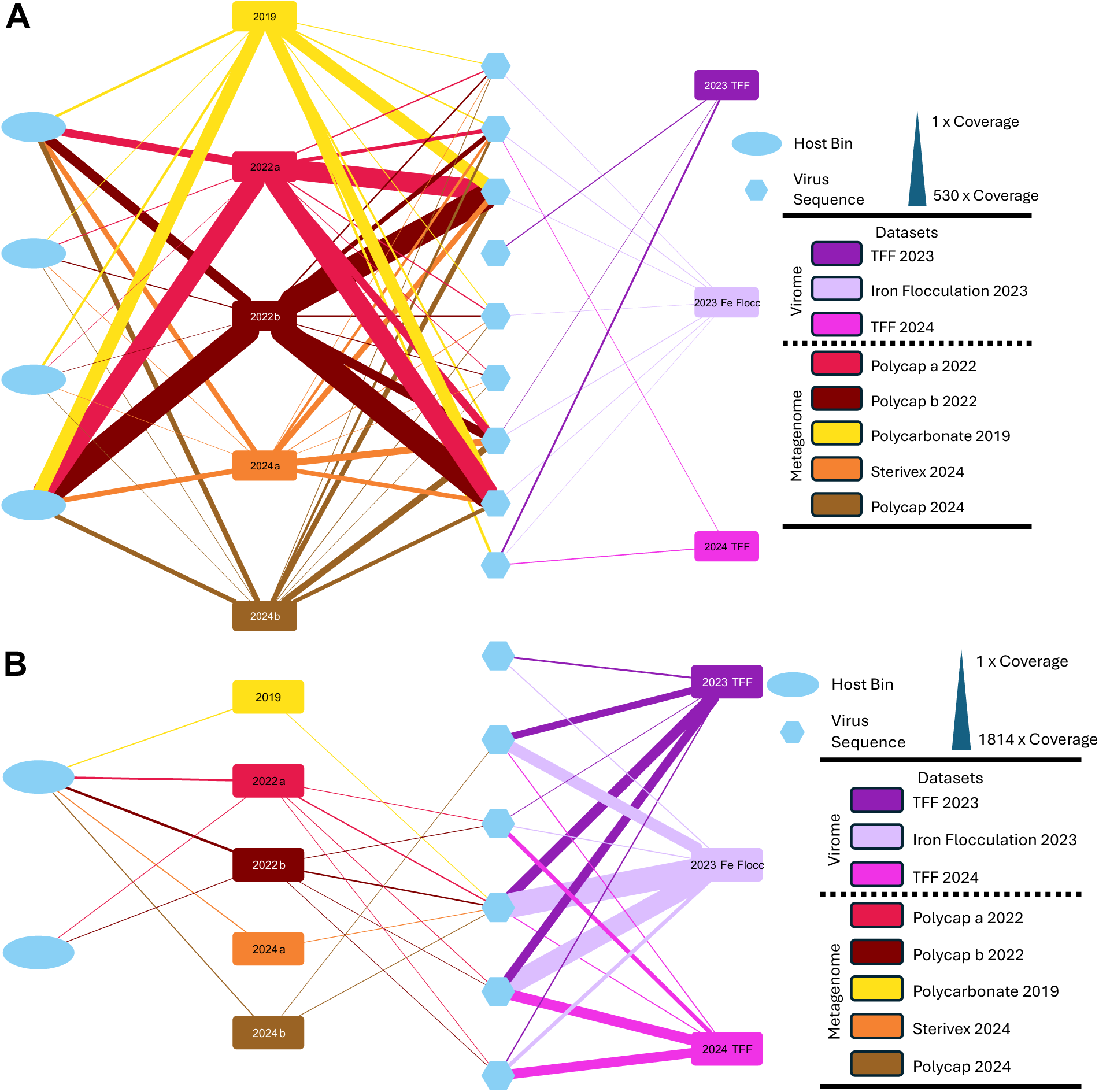
(A) *Rhodocyclaceae* bins and associated viral sequences connected by coverage from different datasets. Ovals represent two *Rhodocyclaceae* bins and hexagons represent the associated viral sequences. Coverage from each of the datasets (rectangles) is represented by the width of the edges. (B) *Thiomicrospiraceae* MAG and viral sequences linked to the samples where there is an average of 1x coverage and higher. Ovals represent two *Thiomicrospiraceae* bins and hexagons represent the associated viral sequences. Coverage from each of the datasets (rectangles) is represented by the width of the edges.

## Discussion

### The virus-like particle (VLP)-to-cell ratio is lower than or similar to other subsurface environments

Our hypothesis of a low (<10) VPR/cell ratio was supported by the data. Cellular counts agree with previous research on this aquifer, in both abundance (∼10^6^ cells/mL) and cellular morphologies [32]. The VLP:cell ratio of ∼1 is on the low end of a range of values reported for other groundwater environments [12, 19–20, 22, 61] and lower than the ratios in marine sediments [63]. Reported ratios ranged from 1.1 to 8.1 for a Colorado aquifer, 1.1 to 18 for groundwater accessed via the Äspö Hard Rock Laboratory, and 0.13 to 2.44 [22] and 0.4 to 6.1 [20] for an Australian aquifer. Parikka et al. compiled an average VLP:cell ratio of 5.9 from several studies published to that date, which is lower than those calculated for marine, freshwater, thermal spring, and soil environments [61]. The authors attribute the low groundwater ratios to the oligotrophic nature of aquifers [61]. While depth appears not to be a factor affecting VLP:cell ratios in previous subsurface studies [20–22], none of these was at as great a depth as the 1200-mbls aquifer considered here.

### Most viruses in the aquifer are novel

The second hypothesis, that novel viruses would be present in the aquifer, was strongly supported. The viral community within the fracture waters contains many unique viruses, with only 1,003 out of 4,499 clustering with at least one cultured virus and 1,320 out of 4,499 clustering with environmental viruses from similar systems. With only 22% of viral sequences clustering with a reference virus, the majority of the viruses present can be considered novel. Of the ones that cluster tightly enough to infer taxonomy by shared gene content, the vast majority of the sequences were classified to *Caudoviricetes*. These are tailed dsDNA bacterial and archaeal viruses that are the most abundant viruses on Earth [64]. While all the viruses identified here exist as dsDNA in the capsid (*Caudoviricetes* and *Tectiliviricetes*) or when in a persistent state (*Faserviricetes*), methodological constraints exclude most ssDNA and all RNA viruses; however, a growing body of knowledge highlights the importance of RNA viruses globally [65]. While a few ssDNA viruses were identified, other ssDNA viruses belonging to the family *Microviridae* were excluded due to the ΦX174 sequencing control used.

### The most common viruses are associated with abundant and also rare hosts

The third hypothesis was at least partially supported in that the majority (83%) of the microbial MAGs from the fracture water appear to have at least one associated viral sequence as seen through host prediction. This included MAGs representing abundant microbes such as those belonging to *Rhodocyclaceae* but also MAGs representing some rarer bins like *Thiomicrospiraceae* and *Moraxellaceae*. Those MAGs without any viruses predicted to infect them range from common to rare, but many of the hosts without associated viruses have much lower coverage. While abundant hosts were targets for some viral sequences, they were not the only targets, meaning the hypothesis is not fully supported.

The microbial community in the fracture water is dominated by *Pseudomonadota*, with *Alphaproteobacteria* and *Gammaproteobacteria* also relatively abundant in 16S rRNA gene libraries and MAGs. The most abundant OTU homologs are classified as *Rhodocyclaceae*, which are also among the most abundant MAGs. The three *Rhodocyclaceae* bins encode the ability to metabolize thiosulfate with a complete Sox system in one bin along with another bin containing *dsr*. The bin containing a complete Sox system also has all genes needed for denitrification. A previously published survey of the microbial community at Moab Khotsong 1200 level included MAGs for members of the families *Rhodocyclaceae*, *Thiobacillaceae*, *Rhodobacteraceae*, *Burkholderiaceae*, *Desulfurivibrionaceae*, *Thiomicrospiraceae*, LZORAL124-64-63, *Halothiobacillaceae*, *Micavibrionaceae*, *Magnetospirillaceae*, *Lentimicrobiaceae*, UBA5603, and UBA918), all of which are represented in bins reconstructed in the present study. The dominance of *Pseudomondata* over *Bacillota* agrees with other relatively young fracture waters [5]. Microbes that commonly dominate subsurface communities in deeper, older, and completely anoxic fracture waters such as methanogens and acetogens [5] appear not to be present here.

To evaluate host-virus associations in the aquifer, we compared the relative abundance of viruses and their predicted hosts in the <0.2 μm and >0.2 μm fractions across all years. First, to compare the viral sequences associated with the most abundant hosts, we examined viruses associated with *Rhodocyclaceae* MAGs. A coverage map was made by connecting the host MAGs and their associated viruses to each of the eight datasets based on the coverage (Fig. 4A). While viruses are present in datasets from the <0.2 μm and >0.2 μm fractions, coverage of the *Rhodocyclaceae*-associated viruses is much higher (and consistently high) in datasets from the >0.2 μm fraction. Three of the viral contigs associated with the *Rhodocyclaceae* bins include genes related to integration, which, along with the high coverage of these contigs in the >0.2 μm fraction (Fig. S8), suggests that these sequences are mainly integrated into the host genomes.

In contrast, there were few hosts for which the associated viral sequences had higher coverage in the <0.2 µm fraction; one of these was *Thiomicrospiraceae* (Fig. S9). A similar coverage map shows that, in contrast to *Rhodocyclaceae*, viral sequences associated with *Thiomicrospiraceae* are consistently more abundant in the datasets from the <0.2 μm fraction, whereas they are present but rare in the >0.2 μm fraction. This indicates that the *Thiomicrospiraceae* viruses existed as free virions at the time of sampling, and were therefore generated via lysis. However, viruses predicted to infect *Thiomicrospiraceae* do have genes related to lysogeny, suggesting that temperate phages that infect this dominant group of bacteria can shift between killing and piggybacking the winner.

Similarities between this study and the virome study at the Äspö hard rock laboratory include dominance of *Pseudomonadota* hosts and also of viruses infecting members of the candidate phyla radiation, now termed *Patescibacteria*, a group of divergent microorganisms first identified in an aquifer near Rifle, Colorado in the U.S. [67]. Members of the *Patescibacteria* are small (≤0.2 µm diameter), have limited metabolic capabilities, and depend upon other bacteria or archaea [68]. While few members exist in culture [69], they have been found to play an important role in biogeochemical cycling in some terrestrial subsurface sites [70–73]. Fewer *Patescibacteria*-associated viral sequences were found here than at Äspö, but the *Patescibacteria* host cells and their associated viruses could have passed through the 0.2-µm filter. In general of the dominance of *Pseudomonadota* viruses and the presence of *Patescibacteria* viruses are similar to findings at Äspö and other subsurface studies [24, 25, 27].

### Aquifer viruses contain a variety of AMGs

While many of the viral genomes in this study harbor AMGs, most are focused on the metabolisms of cofactors, vitamins, amino acids, and nucleotides. Some AMGs such as CysH are involved in nutrient cycling and can potentially influence the geochemical cycling of sulfur [74]. The majority of identified AMGs belong to Class I AMGs, i.e. genes that encode metabolic functions involved in energy, amino acid, lipid, carbohydrate, or cofactors/vitamins [13, 14]. These types of AMGs are typically associated with a lytic lifestyle [15]. Other AMGs such as those involved in folding, sorting, degradation, peptidases, transporters, metal resistance, or xenobiotics degradation are Class II AMGs. While Class II AMGs are present in both lytic and lysogenic viruses, some are especially associated with lysogeny; these include genes for membrane transport, chaperone synthesis or prokaryotic defense mechanisms [15]. While each class is approximately equally represented in the community-wide AMG profile, slightly more appear to be associated with Class II AMGs and, by extension, temperate viruses.

In addition to the AMGs described earlier (Figs. 4 and Fig. S10), two medium-quality viral contigs according to CheckV contained 5-6 AMGS (Fig. S10). The first, NODE_1060, encodes 6 AMGs, including a *mec*, [CysO sulfur-carrier protein]-S-L-cysteine hydrolase, *metK*; S-adenosylmethionine synthetase, and the four major genes for the synthesis of the precursor of 7-deazaguanine (a GTP cyclohydrolase, *queC*, *queD*, and *queE*) [75]. *mec* is involved in the sulfur relay system and assimilatory sulfur reduction pathways [76]. As a functional gene for viruses, this may be involved in redirecting host metabolism towards cysteine biosynthesis [77]. This is supported by the presence of *metK*, also a part of cysteine metabolism by degrading methionine and producing sulfide and S-adenosylmethionine [78]. The genes involved in the synthesis of the precursor of 7-dezaguanine, 7-cyano-7-deazaguanine, are important as a DNA modification system for avoiding host defense mechanisms such as CRISPR-Cas or restriction modification systems. The second virus genome fragment encodes 5 AMGs including a Concanavalin A-like lectin; extracellular arabinose, *phoH*; phosphate starvation-inducible protein, *nrdA*/*nrdB*; ribonucleotide reductase (RNR), dUTP pyrophosphatase, and *mec*; [CysO sulfur-carrier protein]-S-L-cysteine hydrolase. Concanavalin A-like lectin, extracellular arabinose may be a domain of a tail fiber protein related to the binding of the virion to the host [79]. *phoH* is a widely abundant AMG in viruses infecting a wide host range; its function remains unknown other than its relationship to phosphate metabolism [78, 80, 81]. Both *nrdA*/*nrdB* and the dUTP pyrophosphotase are involved in the regulation of deoxyribonucleotides where *nrdA*/*nrdB* are involved in the biosynthesis of deoxyribonucleotide triphosphates [82] while dUTP pyrophosphotase is involved in the cleavage of dUTP to prevent its incorporation into newly synthesized DNA [83].

### Viral communities and viral-microbial interactions appear to be temporally stable and suggest a “piggyback-the-persistent” model

The presence of lysogeny-related genes encoding integrases and excision enzymes in the dominant bacteria, combined with low VLP:cell ratios, suggest that lysogeny is the dominant lifestyle among viruses in the deep aquifer. This could be consistent with “piggyback-the-winner” or “piggyback-the-persistent” interactions. The kill-the-winner model associates lytic viruses with fast-growing hosts and has been modeled using Lotka-Volterra equations for fluctuating populations, with the peaks in populations of viral predators following peaks in cellular prey [9, 33]; such fluctuations are not evident in either the relative abundances of bacterial or viral sequences observed here. The stability of the microbial and viral communities gives a strong indication of lysogeny over lysis and is also likely an indication of stable geochemical conditions. Dominance of lysogeny has been noted in oligotrophic ocean water [84], and the concept of piggyback-the-persistent was developed for a low-nutrient, oligotrophic aquifer with similar cell abundance (∼10^6^ cells/ml) and low VLP:cell ratios (≤0.6) [12] as those seen here, which are typical of groundwater environments, including this dolomitic aquifer. Dissolved organic carbon in this aquifer has been reported at 3.4 mg/L, with only 0.042% of this comprising easily metabolized low molecular weight acids [31]. The 1-4 mg/L sulfide, along with 0.074 mg/L dissolved oxygen, provide energy for chemolithoautotrophc sulfur-oxidizers such as the *Thiomicrospiraceae*, despite this being a low-energy, oligotrophic environment overall. Considering the stability of communities and the generally stressful nature of life in a low-energy environment, implementation of a piggyback-the-persisters approach appears to be a beneficial strategy for viruses in stable groundwater aquifers that are abundant in the deep terrestrial subsurface.

### Recommendation for metaviromic analysis of deep groundwater

Viral sequences are commonly found within metagenomic data generated from environmental samples and should probably be given greater consideration. However, the present study shows, at least for groundwater studies using a filtration step, that the viruses caught on the filter along with the cells are mostly different from those that pass through the filter. These could include prophages and larger free virions, although none of these sequences were related to giant viruses [85]. Inclusion of both approaches may give the best overall picture of the virome. Comparison of the two methods for the concentration of free viral particles (iron flocculation and TFF) showed only minor differences in the viral community; however, many of the most abundant free viral particles were recovered by both methods. Iron flocculation has the advantage of requiring less equipment but may yield fewer viruses; two additional iron flocculation samples were prepared for this study but were not successfully sequenced, whereas TFF recovered enough viruses from the same samples for sequencing. Other investigators have found TFF to be superior in various surface water environments [86, 87] and so the extra investment is likely warranted.

## Conclusions

The deep (1200 mbls) aquifer studied here contains diverse viruses, most of them novel. VLP:cell ratios are low (∼1), consistent with but lower than the ratios for most previously studied, shallower aquifers. The majority of the bacterial families (83%), including the dominant *Thiomicrospiraceae* and *Rhodocyclaceae*, have potential viruses. Many of the AMGs are type II, favoring persistence of the host rather than lysis. The low VLP:cell ratios, the association of dominant microbes with viruses, the presence of lysogeny-related genes in viruses associated with the dominant bacteria, the type II AMGs, and the stable cell and virus community compositions combined with the oligotrophic nature of the aquifer strongly suggest piggyback-the-persistent interaction.

## Acknowledgements

We thank Harmony Gold Mining Company, Ltd. for access to Moab Khotsong Mine and we thank Bennie Liebenberg for serving as liaison to Harmony and for assistance underground. We thank Emil Ruff and Jens Kallmeyer for help in sampling.

## Author contributions

N.E.J. and T.L.K. conceived and designed the research and carried out field sampling. T.L.K and T.C.O. obtained funding, N.E.J., C.S., D.S.J carried out the analyses. N.E.J, T.L.K., and D.S.J. wrote the initial manuscript. C.H., D.N., and C.S. edited the manuscript.

## Data availability

Data are provided within the manuscript or supplementary information. Raw sequence data are available under BioProject accession number PRJNA1348709. High confidence viral sequences from the co-assembly are available at doi.org/10.6084/m9.figshare.30443120. Metagenome assembled genomes from the >0.2 µm coassembly are available at doi.org/10.6084/m9.figshare.30445388.

