## Supplementary methods text for "Diverse but temporally stable virome in a 1.2 km deep karst aquifer accessed via Moab Khotsong mine, South Africa"

Short title: Diverse stable virome in a deep aquifer

Nathaniel E. Jobe[1], Daniel S. Jones[2,3] (https://orcid.org/0000-0003-4556-0418), Julio César Castillo Hernandez[4], Devan Nisson[5], Cassandra H. Skaar[1, 2], Tullis C. Onstott[6], Thomas L. Kieft[1], (https://orcid.org/0000-0003-4350-9416)

Author for correspondence: Thomas L. Kieft, Department of Biology, New Mexico Institute of Mining and Technology, Socorro, NM 87801 U.S.A.


[1]Department of Biology, New Mexico Institute of Mining and Technology, Socorro, NM 87801 U.S.A.

[2]Department of Earth and Environmental Science, New Mexico Institute of Mining and Technology, Socorro, NM 87801 U.S.A.

[3]National Cave and Karst Research Institute, Carlsbad, NM 88220 U.S.A.

[4]Department of Integrated Science, Area of Cell Biology, University of Huelva, Huelva, Spain

[5]NASA Postdoctoral Program, Ames Research Center, Moffett Field, CA, 94035 U.S.A.

[6]Department of Geosciences, Princeton University, Princeton, NJ 08540 U.S.A.

### 16S rRNA gene amplicon libraries

rRNA gene libraries were prepared from DNA extracted from filters (representing the microbial or >0.2 µm fraction) by amplifying the V4 hypervariable region of the 16S rRNA gene using 515f/806r primers [1]: 515f modified GTG YCA GCM GCC GCG GTA A; 806r modified, GGA CTA CNV GGG TWT CTA AT) and Nextera adaptors (forward tail, TCG TCG GCA GCG TCA GAT GTG TAT AAG AGA CAG; reverse tail, GTC TCG TGG GCT CGG AGA TGT GTA TAA GAG ACA G). PCR conditions were 5 minutes at 95 °C, followed by 30 cycles of 95 °C for 45 s, 50 °C for 60 s, and 72 °C for 90 s. Libraries were barcoded, pooled, and sequenced on an Illumina MiSeq platform or Element Biosciences AVITI (paired end, 2 × 300 bp) at the University of Minnesota Genomics Center.

Forward reads were trimmed to an average quality of ≥28 with sickle v1.33 [2] and residual adaptors were trimmed with CutAdapt v2.6 [3]. Forward and reverse reads were assembled using PEAR [4] and operational taxonomic unit (OTU) calling at 97% identify and chimera removal was performed using the UPARSE pipeline [5]. Taxonomy of representative sequences for each OTU was determined against the Silva database v138 [6] using mothur v1.36.1 [7]. A summary of extraction kit, sequencing platform, and quality-controlled reads are summarized in Supplemental Table 1.

Statistical analyses were performed in R v4.4.1 [8] using the vegan package v2.7-0 [9]. Raw OTU counts were standardized using the Hellinger transformation by converting to proportion and performing a square root transformation. Hierarchical agglomerative cluster analysis was performed using Bray-Curtis distance and UPGMA (unweighted pair group method with arithmetic mean) linkage. For two-way cluster analysis, libraries were clustered (R-mode clustering) based on the 50 most abundant OTUs.

### Metagenome analysis

The four metagenomic datasets from the filters, representing the >0.2 μm fractions, were trimmed using Sickle v1.33 [2] to a minimum quality score of 28 and a minimum length of 50 base pairs. Remaining Nextera adaptor sequences on the 3' ends of the reads were removed with Cutadapt v2.6 [3]. An additional metagenomic dataset downloaded from the Sequence Read Archive (SRA; accession: SRX19717538 [10]) was also treated to the above quality control measures. Initial assembly on each of the five datasets to check general assembly statistics was performed with metaSPAdes v4.0.0 [11] and the assembly statistics were verified using the assemblethon_stats.pl script [12]. Then the five datasets were concatenated and co-assembled using metaSPADes v4.0.0 [11] using default options. Reads from each dataset were mapped to the coassembly using bowtie2 v2.5.4 [13] and SAMtools v1.9 [14]. Coverage of each contig was estimated using a modified version of the jgi_summarize_bam_contig_depths script available as part of MetaBat2 [15].

Using the mapped reads from the metagenome datasets, bins were generated with MetaBat2 v2.17 [15] and were quality checked with CheckM v1.2.3 [16] and CheckM2 v1.1.0 [17]. Bins were taxonomically classified using gtdb-tk v2.4.0 [18] and were manually refined with anvi'o v8 [19, 20]. High-quality bins (>50% completion and <5% contamination) were added to the iPHoP v1.3.3 [21] database to improve host prediction. The multiple sequence alignment produced as part of the de_novo_wf of gtdb-tk of the 122 bacterial marker genes from the bins was used to make a tree using FastTree v2.1.11 using the JTT+CAT model to infer approximately-maximum-likelihood phylogenetic trees [22] and visualized using the Interactive Tree of Life (iTOL) v7 [23]. The average coverage of the bins was calculated from the depths file produced by MetaBat2. Overall metabolic capabilities of the high-quality bins were analyzed using DRAM v1.5.0 [24].

The metagenomic datasets prepared from the <0.2 μm fraction using iron flocculation and TFF were quality filtered and trimmed as for the metagenomes from the >0.2 μm fraction described above. Low-quality reads were trimmed using Sickle v1.33 [2] and remaining adapter sequences were removed from the 3' end using Cutadapt v2.6 [2]. As above, the three virome datasets were assembled individually with metaSPAdes v4.0.0 to verify library quality. All datasets (3 metaviromes and 5 metagenomes, including the metagenome from Nisson *et al.* [10]) were concatenated. The concatenated reads were used to coassemble the datasets. Reads were assembled using metaSPAdes v4.0.0 [11] using default parameters and the assembly statistics were verified using the assemblethon_stats.pl script [12]. Reads from the individual datasets were then mapped to the coassembly to determine the coverage from each of the datasets. Mapping was done using bowtie2 v13.3.0 [13] and SAMtools 1.22 [14]. Coverage was determined using the modified version of the jgi_summarize_bam_contig_depths script available as part of Metabat2 v2:2.18 [15] as above.

Viral sequences were identified using VirSorter2 v2.2.4 [25], DeepVirFinder v1.0 [26], and VIBRANT v1.2.1 [27], based on the method of Muscatt et al. [28]. Contigs were classified as viral if they had a VirSorter2 score greater than 0.5, a deepvirfinder score greater than 0.8 or classified as viral by VIBRANT. Programs were run with default settings, specifying a 2 kb minimum length. Viral contigs were proofread using CheckV v1.0.3 [29] and DRAM-v v1.5.0 [26] based on the VirSorter2 (VS2) standard operating procedure (SOP) [30].

The VirSorter2 SOP separates the viral contigs into four categories; keep_1, keep_2, manual_check, and discard. The only change from the SOP was to include criteria from DeepVirFinder (DVF) and VIBRANT in the keep_2 criteria. To be placed into keep_1, the viral contig only needed a minimum of one viral gene according to CheckV. For keep_2, the VS2 SOP requires no viral genes plus one of the following: no host genes, a VS2 score greater than 0.95 or more than two hallmark genes according to VS2. In addition, the criteria from Muscatt et al. [28] for the most confident prediction from DVF and VIBRANT were added. This means the contig could also have a DeepVirFinder score $\geq 0.95$ and $p$-value $\leq 0.05$ or a VIBRANT quality of either "high quality draft" or "complete circular". To be added to manual_check they must not be in either keep category, have no viral genes in CheckV, have only 1 host gene according to CheckV, and be longer than 10 kb. If the contig could not be classified into any of the three categories above, it was discarded. For these, all of keep_1 were kept as high-confidence viral contigs. Keep_2 contigs were screened using the output from DRAM-v looking for annotations related to carbohydrate kinases, glycosyltransferases, nucleotide sugar epimerases, nucleotidyltransferases, plasmid stability, or endonucleases. If the keep_2 contig contained any annotations related to these, it was moved to the manual check category. The final step of proofreading the manual_check category involves screening the contigs for the criteria specified in the VS2 SOP. Additionally, contigs with host genes equaling more than three times the number of viral genes were excluded from downstream analysis. Viral contigs that were retained through proofreading were considered high confidence viral contigs and were then binned using vRhyme v1.1.0 [31] by using the auxiliary script coverage_table_convert.py to convert the output of jgi_summarize_bam_contig_depths to the proper format for vRhyme and using that output to inform the binning process. The best bins were linked using link_bin_sequences.py and concatenated with the unbinned sequences extracted using extract_unbinned_sequences.py.

For the gene sharing network analysis using vConTACT2 v2.6.3 [32, 33], protein sequences were predicted from the high-quality viral sequences using Prodigal with the --meta flag. These protein-coding sequences were run through the vConTACT2 gene2genome script to provide the required gene-to-genome mapping file. Then the corresponding protein and mapping files from the 1Jan2025 INPHARED deposit were concatenated to the files from the high-quality viral sequences. These files were used in vConTACT2 specifying no database and using ClusterONE v1.0 [34].

For taxonomic assignment using vConTACT2 and files from INPHARED, protein sequences were predicted from the high-quality viral sequences using Prodigal with the meta flag. These protein-coding sequences were run through the vConTACT2 gene2genome script to provide the required gene-to-genome mapping file. Then the corresponding protein and mapping files from the 1Jan2025 INPHARED deposit were concatenated to the files from the high-quality viral sequences. These files were used in vConTACT2 specifying no database and using ClusterONE v1.0 [34].

For the additional network created using viral sequences in IMGVR v4.1 [35] from marine aquifers, deep subsurface aquifers, deep subsurface groundwater, and freshwater groundwater, protein sequences from were called as above using Prodigal. The protein-coding sequences were combined with the protein-coding sequences for the high-quality viruses and used to make a gene-to-genome mapping file. This was run as above with the addition of the ProkaryoticViralRefSeq211-Merged database. Both cluster maps were visualized in Cytoscape [36].

Host prediction was performed using iPHoP v1.3.3 and a customized database consisting of the default Aug_2023_pub_rw database with the addition of 71 “high quality” metagenome-assembled genomes (MAGs). Briefly, iPHoP uses a combination of six different methods of virus-host linkage and compiles the most likely prediction from each program. The iPHoP output was filtered to focus on the viruses linked to the “high quality" MAGs.

AMGs were identified using VIBRANT v1.2.1 and DRAM-v v1.5.0 . VIBRANT v1.2.1 and VirSorter2 v2.2.4 were rerun on the high-quality contigs and limited to retain as many viral sequences as possible to extract the annotated AMGs from VIBRANT and prepare the files necessary for DRAM-v v1.5.0 from VirSorter2. The output files from VirSorter2’s --prep_for_DRAM setting were used to run both DRAM-v annotated and DRAM-v distill to identify AMGs.

Pharokka 1.7.3 [37] was used to predict coding sequences (CDS) with Prodigal-gv v0.3.1 [38, 39], tRNAs were predicted with tRNAscan-SE v2.0.12 [40], tmRNAs were predicted with Aragorn v1.2.41 [41], and CRISPRs were predicted with CRT via MinCED v0.4.2 [42]. Functional annotations of each CDS were performed by matching each CDS to the PHROGs [43], VFDB [44] and CARD [45] databases using MMseqs2 v13.45111 [46] and PyHMMER v0.10.12 [47]. Further refinement of the annotations was performed with phold v0.2.0 [48], which uses Foldseek v9.427df8a [49] to search the protein structures against database phage protein structures that were generated by any of several programs, but primarily Colabfold [50]. Plots were generated LoVis4u v0.1.2 [51] for linear and comparative plots.
