## Supplementary figures and tables for "Diverse but temporally stable virome in a 1.2 km deep karst aquifer accessed via Moab Khotsong mine, South Africa"

Short title: Diverse stable virome in a deep aquifer

Nathaniel E. Jobe[1], Daniel S. Jones[2,3] (https://orcid.org/0000-0003-4556-0418), Julio César Castillo Hernandez[4], Devan Nisson[5], Cassandra Skaar[1, 2], Tullis C. Onstott[6], Thomas L. Kieft[1], (https://orcid.org/0000-0003-4350-9416)

Author for correspondence: Thomas L. Kieft, Department of Biology, New Mexico Institute of Mining and Technology, Socorro, NM 87801 U.S.A.


[1]Department of Biology, New Mexico Institute of Mining and Technology, Socorro, NM 87801 U.S.A.

[2]Department of Earth and Environmental Science, New Mexico Institute of Mining and Technology, Socorro, NM 87801 U.S.A.

[3]National Cave and Karst Research Institute, Carlsbad, NM 88220 U.S.A.

[4]Department of Integrated Science, Area of Cell Biology, University of Huelva, Huelva, Spain

[5]NASA Postdoctoral Program, Ames Research Center, Moffett Field, CA, 94035 U.S.A.

[6]Department of Geosciences, Princeton University, Princeton, NJ 08540 U.S.A.

**Table S1** Summary of aquifer geochemistry across three years.

| | Year | | |
|---|---|---|---|
| Parameter | 2022 | 2023 | 2024 |
| Temperature (℃) | 26.40 | 22.40 | 23.60 |
| pH | 7.9 | 8.3 | 8.6 |
| EC (mS/cm) | 1.71 | 1.70 | 1.60 |
| ORP (mV) | -222 | -288 | -276 |
| Salinity (g/L) | 0.86 | 0.87 | 0.81 |
| TDS (mg/L) | 1086 | 1112 | 1029 |
| DO (mg/L) | nm[a] | | 0.074 |
| Chemet kits (mg/L) | | | |
| Sulfide | 4.00 | 2.00 | 1.00 |
| DO (0-1 mg/L kit) | 0.05 | >40 | 3.00 |
| Ferrous Iron | 0.16 | 0.00 | nm |
| Iron Total | 0.22 | 0.00 | nm |
| Alkalinity (mg/L) | | | |
| Alkalinity as $CaCO_3$ | nm | 210 | 216 |
| Bicarbonate ($HCO_3^-$) | nm | 256 | 243 |
| Carbonate ($CO_3^{2-}$) | nm | nd[b] | 10 |
| Hardness (mg $CaCO_3$/L) | nm | 46.9 | 66.5 |
| Major anions (mg/L) | | | |
| Bromide | nm | 1.73 | 1.25 |
| Chloride | nm | 269 | 283 |
| Fluoride | nm | 9.8 | 9.32 |
| Nitrate | nm | nd | nd |

| | | | |
|---|---|---|---|
| Nitrite | nm | nd | nd |
| Orthophosphate | nm | 6.24 | 3.77 |
| Sulfate | nm | 137 | 143 |
| Major cations (mg/L) | | | |
| Calcium | nm | 8.34 | 14.5 |
| Iron | nm | nd | nd |
| Magnesium | nm | 6.32 | 7.35 |
| Manganese | nm | 0.006 | 0.039 |
| Potassium | nm | 2.22 | 2.88 |
| Silicon | nm | 7.01 | 6.6 |
| Sodium | nm | 304 | 306 |
| Strontium | nm | 0.246 | 0.454 |
| ICP-MS (mg/L) | | | |
| Aluminum | nm | 0.0038 | 0.0139 |
| Antimony | nm | nd | nd |
| Arsenic | nm | nd | nd |
| Barium | nm | 0.111 | 0.103 |
| Beryllium | nm | nd | nd |
| Boron | nm | 0.953 | 0.916 |
| Cadmium | nm | nd | nd |
| Chromium | nm | nd | nd |
| Cobalt | nm | 0.0428 | 0.0418 |
| Copper | nm | nd | nd |
| Lead | nm | nd | nd |
| Lithium | nm | 0.087 | 0.088 |

| | | | |
|---|---|---|---|
| Molybdenum | nm | 0.003 | 0.003 |
| Nickel | nm | nd | nd |
| Selenium | nm | nd | nd |
| Thallium | nm | nd | nd |
| Thorium | nm | nd | nd |
| Tin | nm | nd | nd |
| Titanium | nm | nd | nd |
| Uranium | nm | 0.007 | 0.0146 |
| Vanadium | nm | nd | nd |
| Zinc | nm | 0.134 | 0.0157 |

[a]nm = not measured

[b]nd = not detected

**Table S2.** Details of 16S rRNA gene amplicon libraries

| Year | Sample name | Filter type | Extraction kit | Sequencing platform | Number of sequences post QC |
|---|---|---|---|---|---|
| 2022 | Polycap 2022 r1 | Polycap | PowerSoil Pro | Illumina MiSeq | 28169 |
| 2022 | Polycap 2022 r2 | Polycap | PowerSoil Pro | Illumina MiSeq | 57148 |
| 2023 | Polycap 2023 | Polycap | PowerSoil Pro | Illumina MiSeq | 45340 |
| 2024 | Sterivex 2024 r1 | Sterivex | PowerWater | Element Biosciences AVITIT | 748531 |
| 2024 | Polycap 2024 | Polycap | PowerWater | Element Biosciences AVITIT | 495226 |
| 2024 | Sterivex 2024 r2 | Sterivex | PowerWater | Element Biosciences AVITIT | 629055 |

**Table S3.** Assembly statistics of individual metagenomic datasets used in this study.

| | Viromes | | | | Mixed metagenomes (cells and viruses) | | | | |
|---|---|---|---|---|---|---|---|---|---|
| Year | 2023 | | 2024 | | 2019 | 2022 | | 2024 | |
| Sample Name | 100kDa TFF | PC_Fe | 1200L-TFF | FE_clean up | Nisson[a] | BF1 | BF2 | PC_b | Steri_a |
| Sample Type | TFF | Iron Flocculation | TFF | Iron Floc-culation | Polycar bonate | Polycap | Polycap | Polycap | Sterivex |
| Sequen-cing platform | Illumina NovaSeq | Illumina NovaSeq | Element Bio-sciences AVITI | Element Bio-sciences AVITI | Illumina NovaSe q 6000 | Illumina NovaSeq | Illumina NovaSeq | Element Bio-sciences AVITI | Element Bio-sciences AVITI |
| Number of scaffold s x 10^3 | 33.69 | 46.93 | 50.42 | 4.16 | 119.33 | 155.44 | 180.13 | 24.70 | 19.54 |

| | | | | | | | | | |
|---|---|---|---|---|---|---|---|---|---|
| Total size of scaffold s (Mbp) | 60.37 | 100.49 | 100.24 | 5.44 | 201.41 | 243.53 | 279.73 | 84.36 | 77.66 |
| Longest scaffold (kbp) | 270.39 | 705.73 | 572.27 | 85.25 | 864.66 | 733.82 | 549.84 | 610.36 | 588.96 |
| Mean scaffold size (bp) | 1792 | 2142 | 1988 | 1309 | 1711 | 1567 | 1553 | 3416 | 3975 |
| Median scaffold size (bp) | 813 | 841 | 885 | 969 | 761 | 723 | 730 | 878 | 874 |
| N50 (bp) | 3174 | 4836 | 3711 | 1514 | 3046 | 2415 | 2348 | 19231 | 25877 |
| L50 | 2946 | 2589 | 3591 | 1030 | 7640 | 10897 | 14101 | 765 | 654 |
| > 10k | 712 | 1052 | 1253 | 10 | 2182 | 2132 | 2417 | 1439 | 1449 |
| >100kb | 21 | 79 | 38 | 0 | 119 | 172 | 175 | 87 | 75 |

[a]Metagenome dataset downloaded from the SRA from Nisson et al. [32].

**Table S4.** Summary of high quality metagenome-assembled genomes (MAGs)

(Attached as a separate spreadsheet)

**Table S5.** Details of auxiliary metabolic genes (AMGs) classified as "other"

(Attached as a separate spreadsheet)

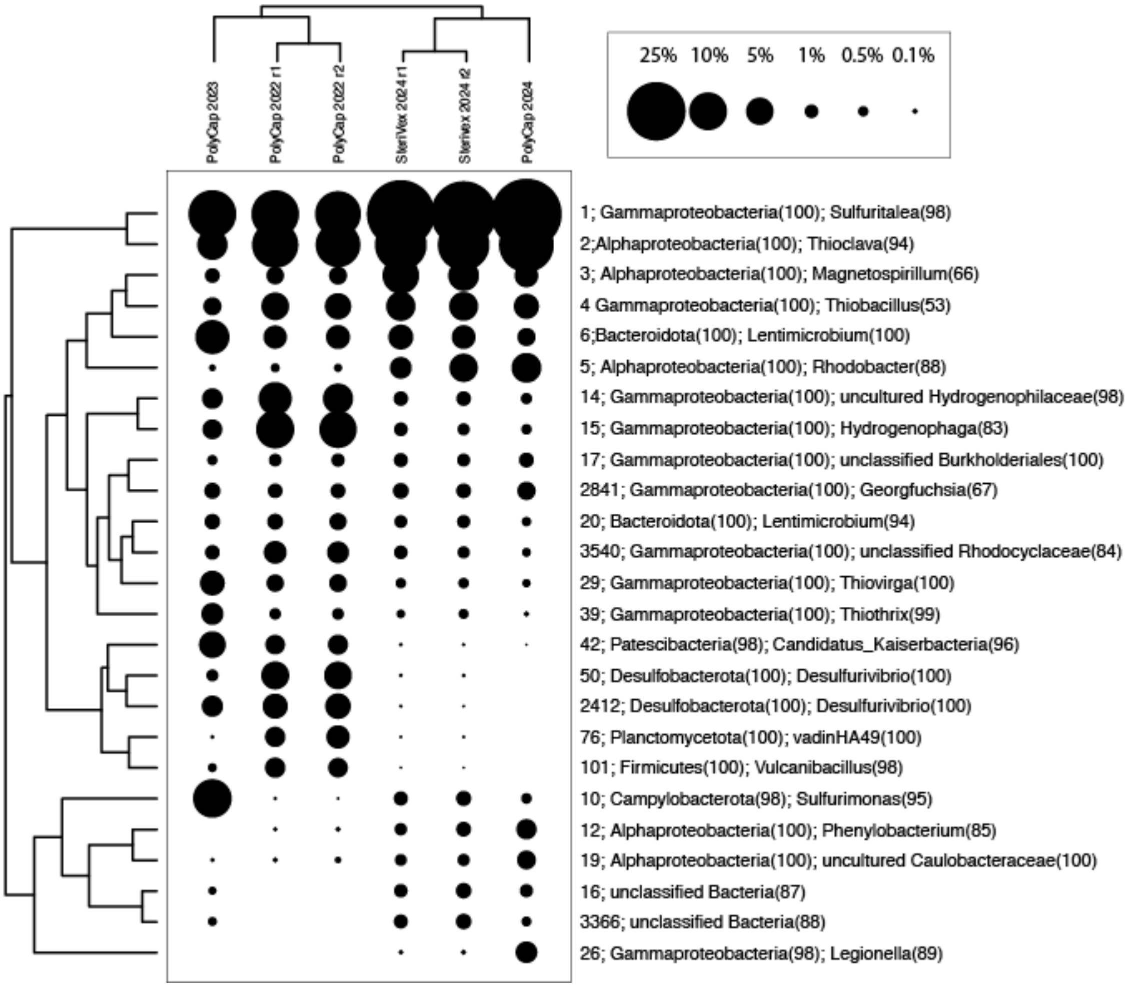


**Figure S1.** Two-way cluster analysis of the twenty-five most abundant OTUs from the 16S rRNA gene amplicon libraries with the associated taxonomy.

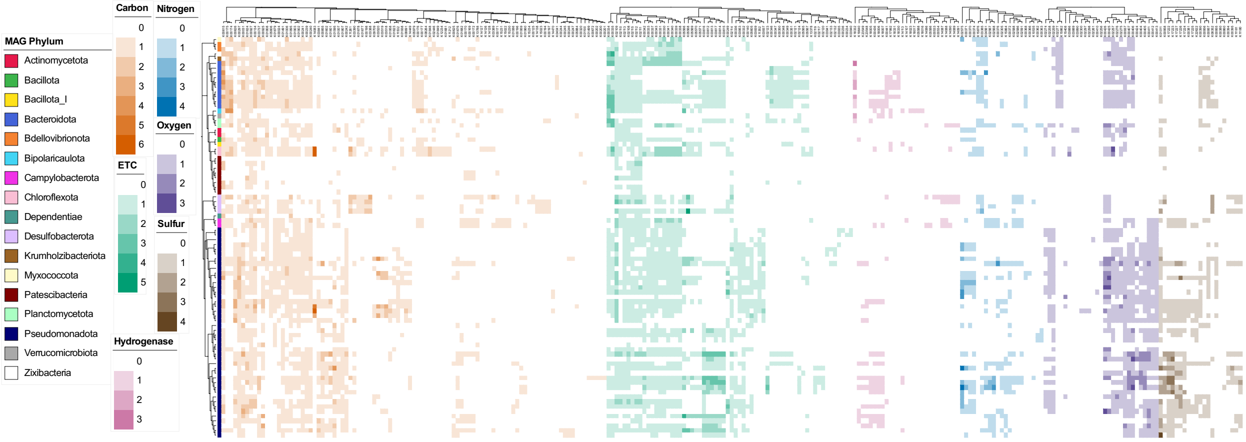


**Figure S2.** Heatmap of additional annotation for the high-quality bins by DRAM for metabolism.

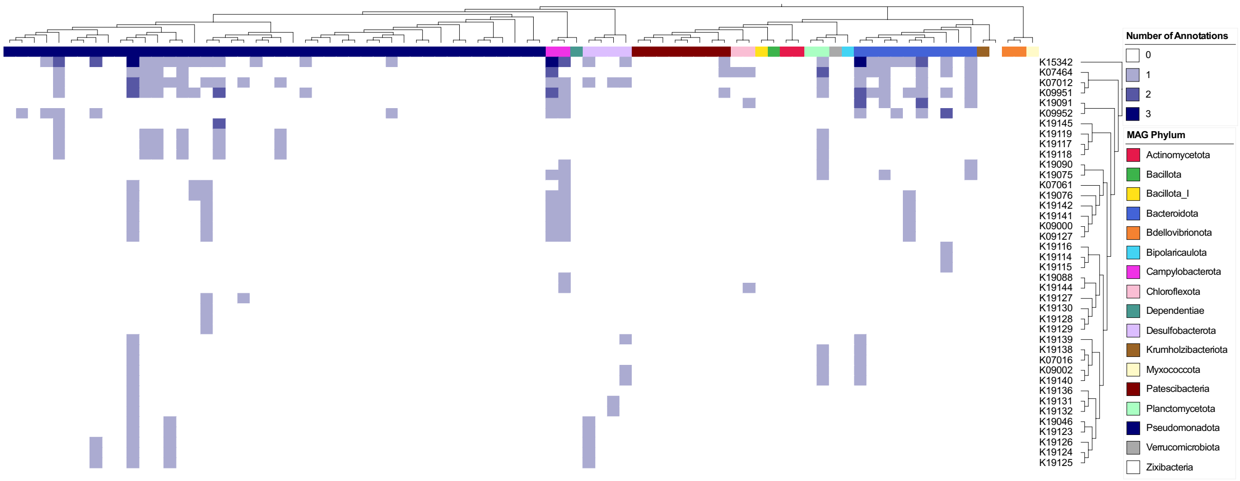


**Figure S3.** Heatmap of CRISPR genes annotated by DRAM for high-quality bins.

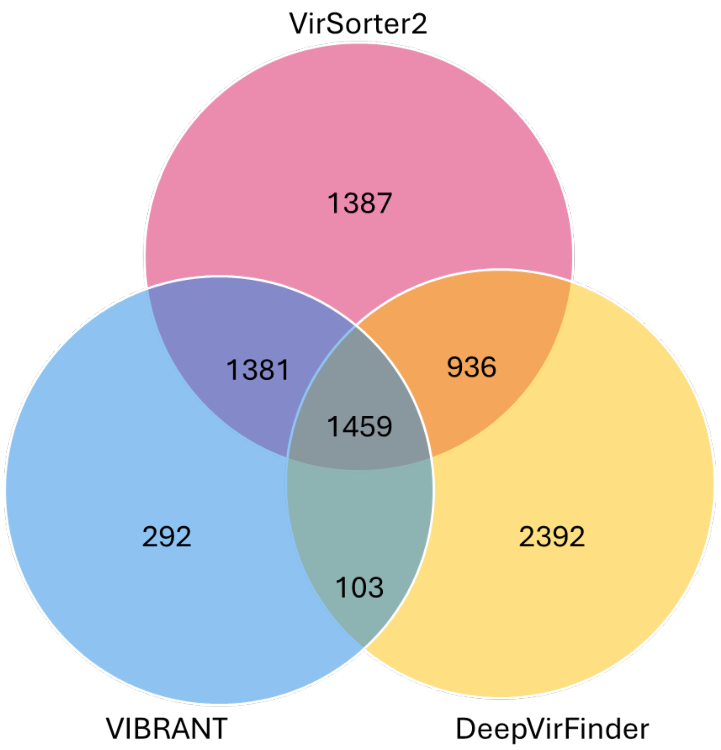


**Figure S4.** Venn diagram of the number of predicted viral contigs and the programs that predicted them before binning the contigs with vRhyme.

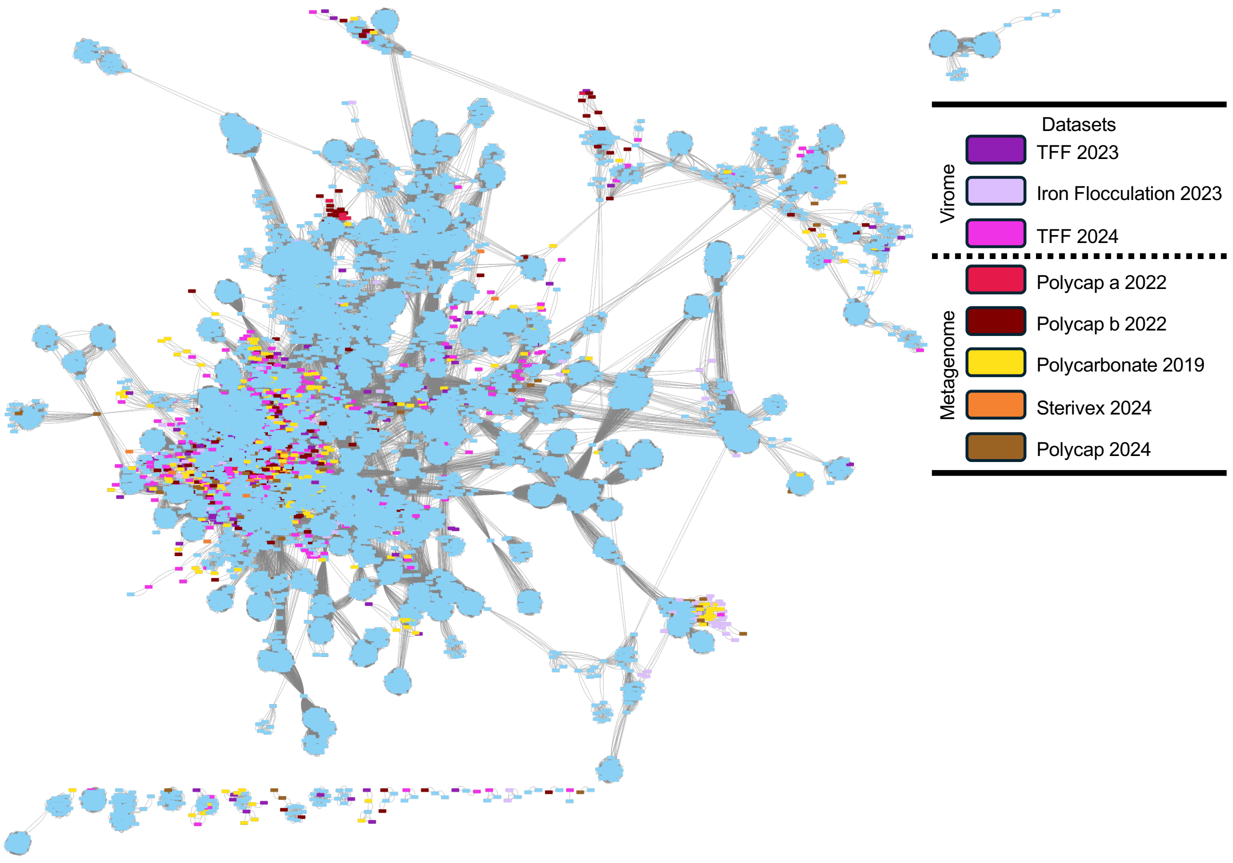


**Figure S5.** Gene sharing network analysis of viral sequences from the coassembly along with with INPHARED database viruses. Sample nodes are colored by the dataset that contributed the most coverage for the sequence, dabase sequences are colored blue. Visualized in Cytoscape.

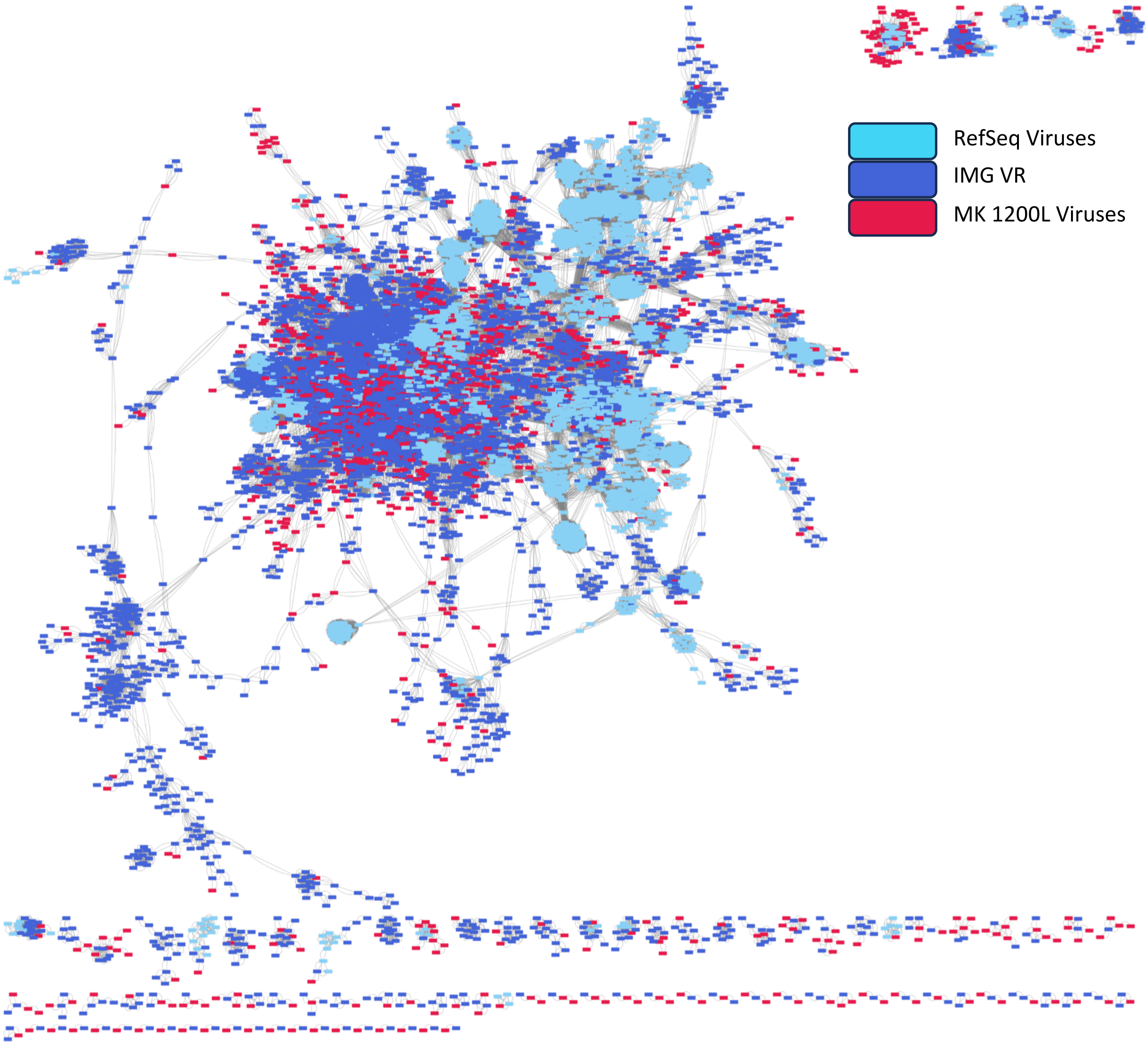


**Figure S6.** Gene-sharing network of viral sequences against environmental viruses from marine aquifers, deep subsurface aquifers, deep subsurface groundwater, and freshwater groundwater from IMG/VR in addition to the RefSeq viral database from vCONTACT2. Visualized in Cytoscape.

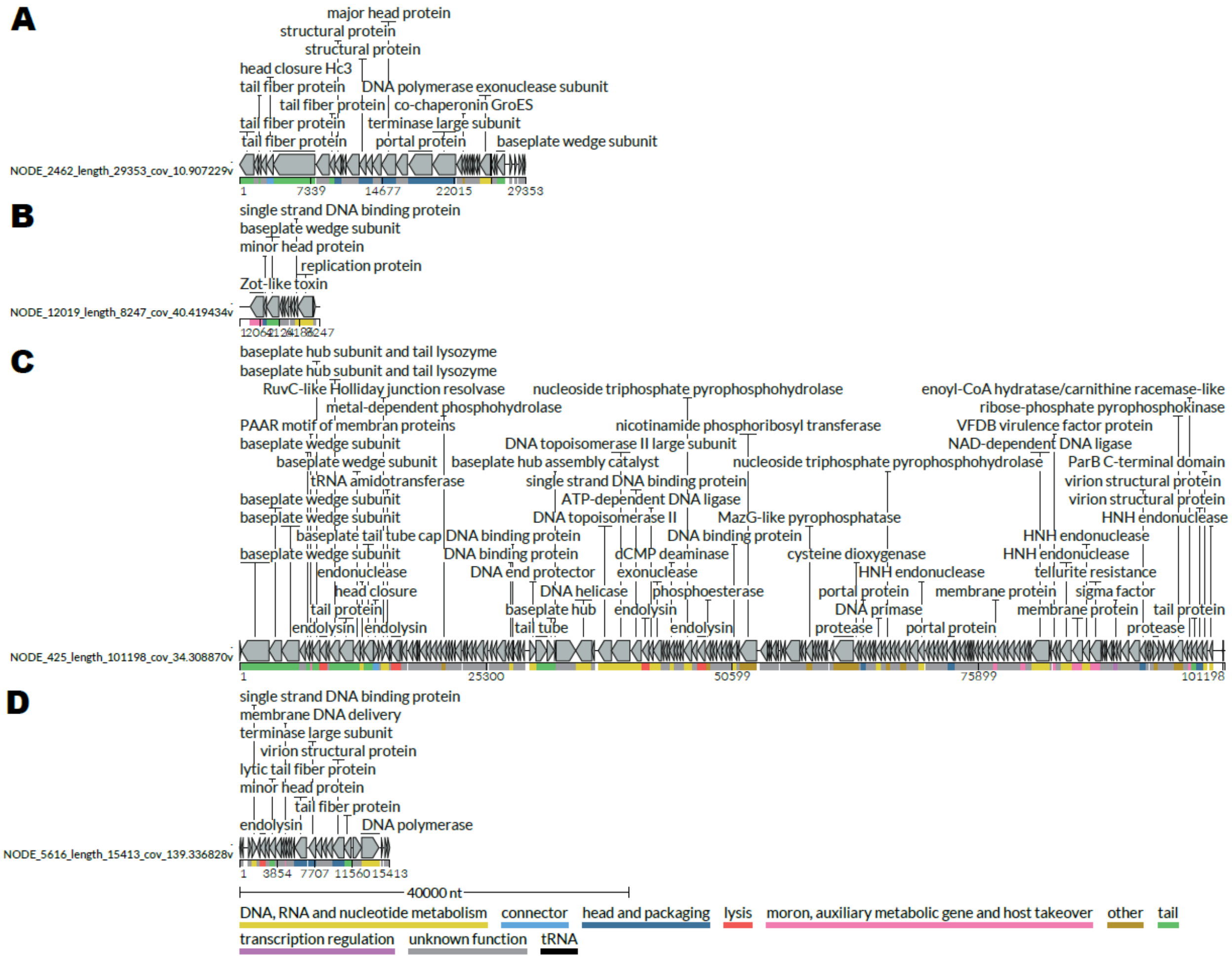


**Figure S7**. Genome maps of viruses that were taxonomically classified to the family level or lower. A) Viral contig classified to the family *Crevaviridae.* B) Viral contig classified to the genus *Inovirus*. C) Viral bin classified to the family *Kyanoviridae.* D) Viral contig classified to the genus *Gammatectivirus*.

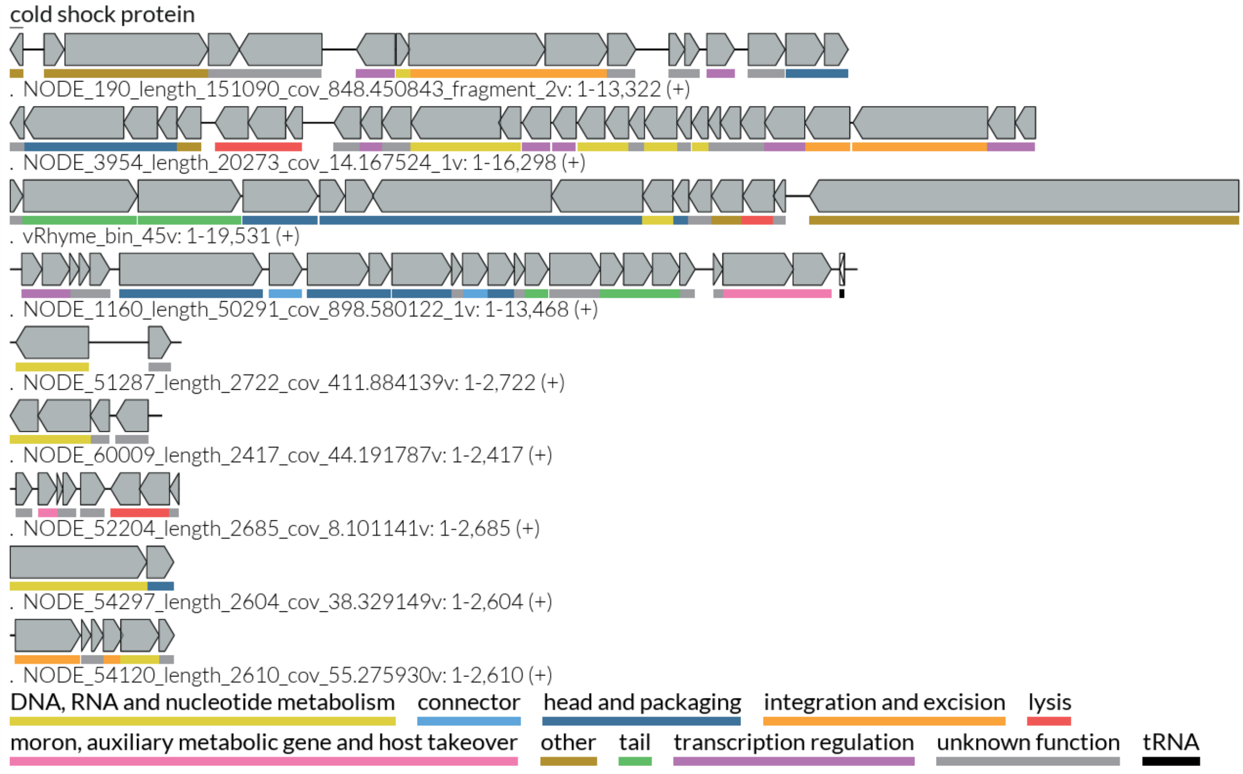


**Figure S8.** Genome maps of the viral sequences predicted to infect the *Rhodocyclaceae* MAGs.

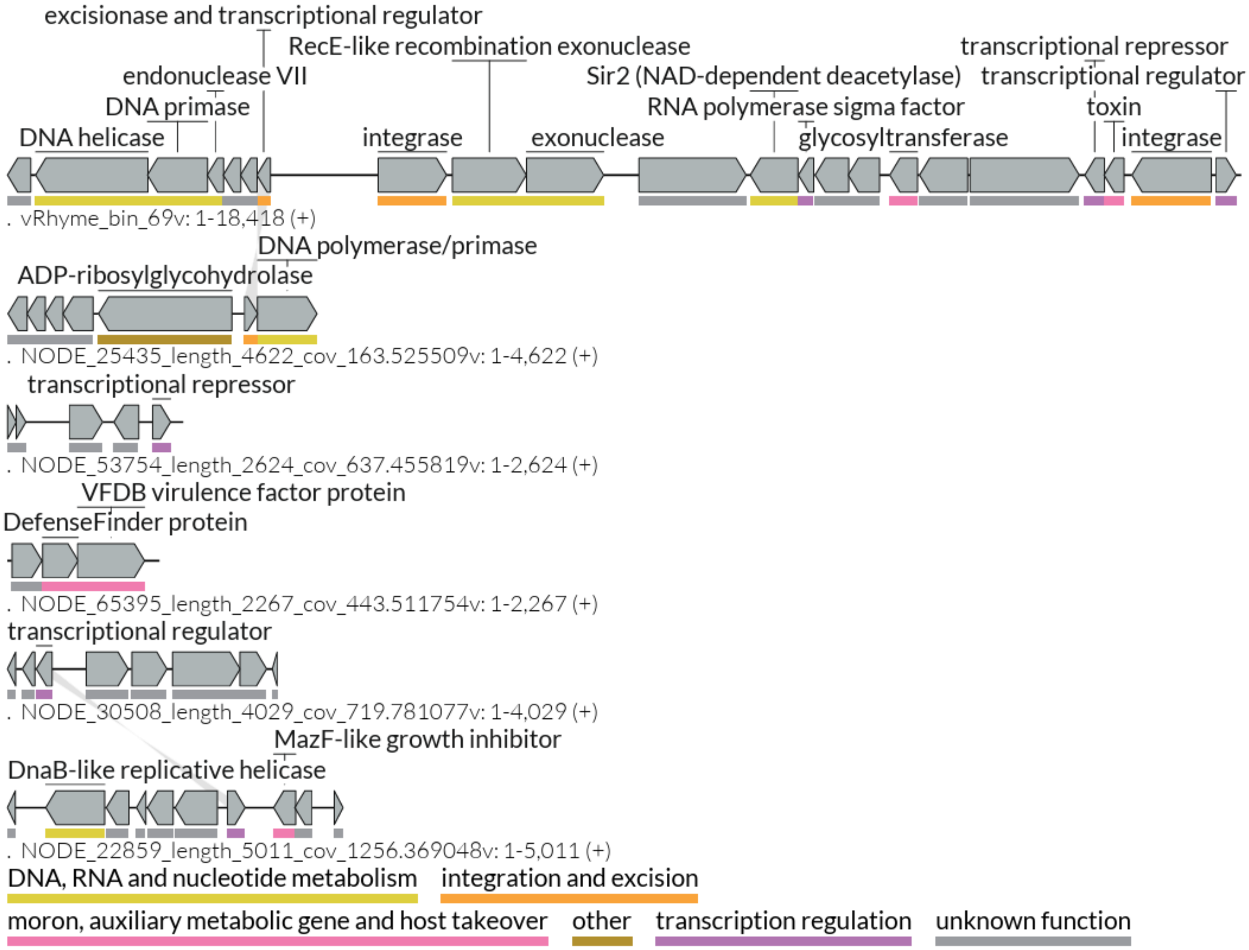


**Figure S9.** Genome maps of the viral sequences predicted to infect the *Thiomicrospiraceae* MAGs. The grey bars between sequences represent protein sequence homology between sequences.

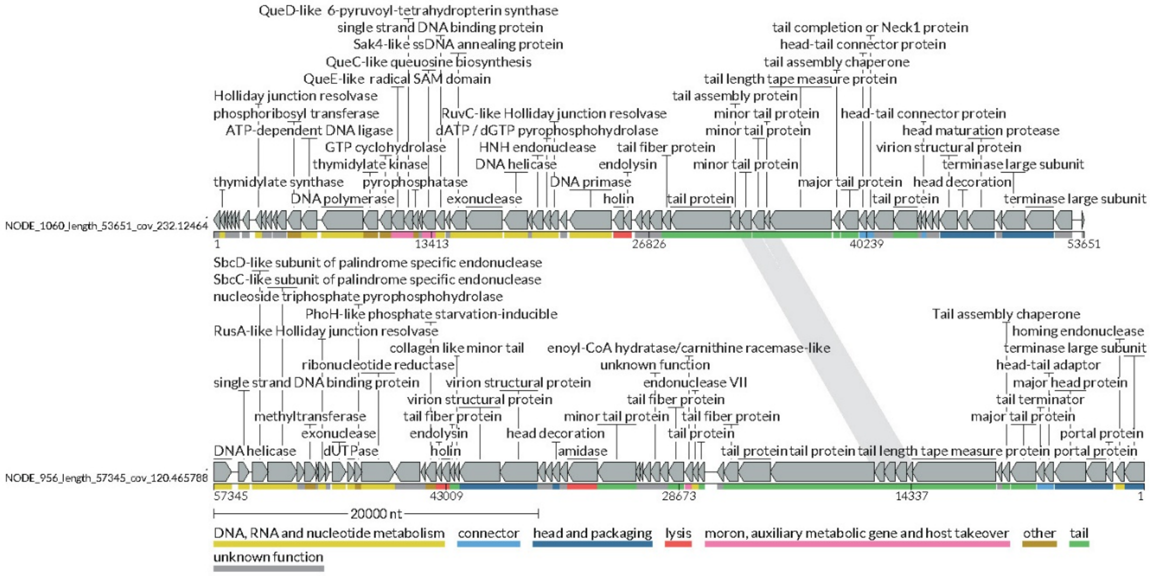


**Figure S10.** Viral contigs containing various AMGs. The grey bars between sequences represent protein sequence homology between sequences.
